# The mechanisms and neural signature of average numerosity perception

**DOI:** 10.1101/2024.04.29.591635

**Authors:** Irene Togoli, Olivier Collignon, Domenica Bueti, Michele Fornaciai

## Abstract

The human brain is endowed with an intuitive sense of number allowing to perceive the approximate quantity of items in a scene, or “numerosity.” This ability is not limited to items distributed in space, but also to events unfolding in time and to the *average* numerosity of dynamic scenes. How the brain computes and represents the average numerosity over time however remains mostly unclear. Here we investigate the mechanisms and electrophysiological (EEG) signature of average numerosity perception. To do so, we used dynamic stimuli composed of 3-12 arrays presented for 50 ms each, and asked participants to judge the average numerosity of the sequence. Our results first show that the weight of different arrays in the sequence in determining the judgement is subject to both primacy and recency effects, depending on the length of the sequence. Moreover, we show systematic perceptual adaptation effects across trials, with the bias on numerical estimates depending on both the average numerosity and length of the preceding stimulus. The EEG results show numerosity-sensitive brain responses starting very early after stimulus onset, and that activity around the offset of the sequence can predict both the accuracy and precision of judgments. Additionally, we show a neural signature of the adaptation effect at around 300 ms, whereby the amplitude of brain responses can predict the strength of the bias. Overall, our findings support the existence of a dedicated, low-level perceptual mechanism involved with the computation of average numerosity, and highlight the processing stages involved with such process.

## INTRODUCTION

Humans and other animals have an innate ability to rapidly estimate the number – or *numerosity* – of objects in a visual scene (e.g., Feigenson et al., 2004). This ability is independent of counting, and produces an approximate estimation prone to errors proportional to the number of items being estimated (e.g., Anobile et al., 2016; but see Testolin & McClelland, 2021). Due to the properties of numerosity perception, like for instance it being subject to perceptual adaptation effects, numerosity has been proposed to represent a “primary” perceptual attribute (Anobile et al., 2016; Burr & Ross, 2008; but see Leibovich et al., 2017 for a different account), that is, one of the fundamental building blocks of our perceptual experience. Research into numerosity perception focused especially on the judgment of items presented simultaneously in space, like arrays of dots. Numerosity however can be computed from several different types of stimuli. For instance, rather than objects distributed in space, numerosity can be extracted from series of events (e.g., brief flashes) presented over time (e.g., Arrighi et al., 2014). Research in this context has shown that different types of numerical stimuli can affect each other via the process of perceptual adaptation (Anobile, Arrighi, et al., 2016; Arrighi et al., 2014) and the serial dependence effect (Fornaciai & Park, 2019a), suggesting the existence of an abstract “number sense” (Anobile, Arrighi, et al., 2016; Arrighi et al., 2014). In terms of neural correlates, numerosity-sensitive brain activity has been observed throughout the visual stream starting from early visual areas, in terms of localisation (Castaldi et al., 2019; DeWind et al., 2019; Harvey et al., 2013; Roggeman et al., 2011), and from very early processing stages, in terms of timing (Fornaciai et al., 2017; Fornaciai & Park, 2018; Park et al., 2016; Temple & Posner, 1998).

While the use of static dot-array stimuli probably remains the most common practice in numerosity perception research, the external environment and the stimuli that our sensory organs receive are rarely static. Perception is indeed a dynamic process, due for instance to the frequent and fast shifts of our gaze and attention to sample the surrounding world. An intriguing question is thus how the visual system computes and process the *average* numerosity of dynamic visual events, involving both the spatial and the temporal dimension. Previous research in this context shows that when presented with dynamic stimuli modulated over time (Togoli et al., 2021) or series of discrete stimuli (Katzin et al., 2021), humans are able to perceive and judge their *average* numerosity with good accuracy and precision. In line with the concept of “summary statistics” (e.g., McDermott et al., 2013; Whitney & Yamanashi Leib, 2018), many studies indeed show that the visual system can easily extract or compute the average value of a feature modulated over time (e.g., Chong & Treisman, 2005; de Fockert & Wolfenstein, 2009; Robitaille & Harris, 2011). In terms of the properties of average numerosity perception, Katzin et al., (2021) showed that the judgment precision of average numerosity tends to increase with the sequence length, leading to better averaging performance when more information is provided. Additionally, Katzin et al.’s (2021) results provided some indications of recency effects, whereby more recent information in the sequence has a larger weight on perceptual decisions, although this effect was observed only in some participants. However, such results were obtained with sequences of relatively long (i.e., 500 ms) discrete stimuli, which likely involve memory rather than perceptual processes.

In the present study, we thus aim to further address the perceptual mechanisms of average numerosity perception, and the neural signature of this process. To do so, we employed a classification task of the average numerosity of dynamic dot-array stimuli (Togoli et al., 2021). Namely, in each trial the participants observed a fast sequence of 3-12 individual arrays varying in numerosity, each presented for 50 ms, and were asked to judge whether their average numerosity was higher or lower compared to a memorised reference. Such a fast dynamic stimulation (i.e., 20 Hz modulation of numerosity) was specifically chosen in order to give the impression of a continuous stimulus, rather than a sequence of static arrays. The average numerosity of the sequence was varied between 15 and 60 dots, and each array in the sequence could vary ± 50% around the mean. Electroencephalography (EEG) was used to assess the brain responses to the stimuli. To understand how average numerosity is computed and represented, we first assessed the weight of different arrays in the sequence (first, middle, last) in determining the judgment. This was done separately according to the different sequence lengths, to test whether increasing the amount of information may affect the temporal weighting profile of the stimuli. Moreover, we assessed perceptual adaptation effects across different trials. With EEG, we measured the neural signature of average numerosity processing and the adaptation effect, and the relationship between behavioural and neural measures, in order to better understand the brain processing stages linked to the representation of average numerosity.

## METHODS

### Participants

The sample tested in this study included 22 adult volunteers (mean age = 23 years, SD = 2.82; age range = 18-31; 1 male). All participants provided written informed consent prior to testing and received monetary compensation for their time (10€/hour). All the participants had normal or corrected-to-normal vision, were naive to the purpose of the experiment, and reported no history of neurological, attentional, or psychiatric disorders. The research protocol was approved by the ethics committee of the International School for Advanced Studies (SISSA) (Protocol 10035-III/13), and was in line with the Declaration of Helsinki. One participant was excluded from data analysis due to equipment failure (i.e., missing electroencephalography data), leaving 21 participants included in the final sample of the study. The sample size of the study was computed with a power analysis based on two previous studies addressing magnitude integration effects (Togoli et al., 2021, 2022). In the power analysis, we conservatively considered the effect of the smallest levels of the interfering magnitude in the Exp. 1a of Togoli et al., 2021, and the effects in Exp. 2 of Togoli et al., 2022. The average effect size (Cohen’s d) computed from these results was d = 0.82. Considering a two-tailed distribution and a power of 95%, the power analysis indicated a sample size of 22 participants.

### Stimuli

The visual stimuli were generated using the routines of the Psychophysics Toolbox (v.3; Kleiner M et al., 2007; Pelli, 1997) in Matlab (r2021b, The Mathworks, Inc.). During the experiment, the stimuli were displayed on a 1092×1080 LCD monitor running at 120 Hz, encompassing a visual angle of about 48×30 degrees from a distance of 57 cm. The stimulus design was based on Togoli et al., 2021, and consisted of dynamically-modulated arrays of dots. Specifically, the dynamic stimuli involved a sequence of multiple briefly-flashed (50-ms each; 20 Hz frequency) dot arrays modulated in average numerosity (i.e., the average amount of dots displayed across all the arrays included in a sequence). The number of dots in each array varied around the mean numerosity of the sequence selected in each trial (±50%). The numerosity of each array was computed before the presentation of the stimulus in order for the sequence to result in a specific average numerosity. The positions of the dots were computed in order to avoid overlapping, considering a minimum inter-dot distance of 2.5 times the radius of an individual dot. Dot sizes and the radius of the area encompassing them were systematically varied in a trial-by-trial fashion, in line with the procedure used in previous studies (DeWind et al., 2015; Park et al., 2016; Fornaciai et al., 2017). The radius of the dots ranged from 6 to 10 pixels, while the radius of the area of the stimulus spanned from 200 to 400 pixel. Each array in a sequence had the same area and the same dot size. The dots were black and white with a 50/50% proportion, and in case of odd numerosities the colour of the exceeding dot was determined randomly. The average numerosity of each stimulus could be either 15, 21, 30, 42, or 60 dots. The number of arrays in the sequence could be 3, 4, 6, 9, or 12 arrays, corresponding to a total duration of the stimulus of 150, 200, 300, 450, 600 ms. The average numerosity range and the number of arrays were combined resulting in a total of 25 different stimulus types. Before the beginning of the session and before each block, we presented a reference stimulus that the participants had to memorise and use as a comparison to provide a judgment. The reference stimulus had the intermediate values of the average numerosity and number of arrays, i.e, it had an average numerosity of 30 dots and was composed of 6 arrays (duration = 300 ms).

### Procedure

The experiment was conducted in a sound-attenuated room, with each participant sitting in front of the computer screen at a distance of about 57 cm. The study involved a classification task of the average numerosity of the dynamic stimuli. Electroencephalography (EEG) was also recorded throughout the session to measure the brain responses to the stimuli. At the beginning of the session, participants were shown the reference stimulus (average numerosity of 30 dots, 6 arrays) that they were instructed to use in order to classify the stimuli in the main task sequence. The reference was displayed ten times. During the task, participants kept their gaze on a central fixation point, and the dynamic stimuli were presented at the centre of the screen. Following the offset of each stimulus, there was a 600-ms interval after which the fixation cross turned red, signalling to the participant to provide a response. The participant was then asked to judge, by pressing the left or the right arrow on the keyboard, whether the average numerosity was lower or higher compared to the memorised reference stimulus (respectively). The time available to provide a response was limited to 1200 ms. If they could not respond within this interval, the next trial started automatically. The inter-trial interval (ITI) was 1100-1300 ms). The trials in which participants were not able to provide a response were excluded from data analysis (1.1% ± 1.2%). Participants received no feedback about their response. The reference stimulus was presented again to the participants before the beginning of each block (displayed five times). Each participant completed a total of 10 blocks of 100 trials, for a total of 1000 trials and 40 repetitions of each combination of average numerosity and number of arrays. Before the start of the session, subjects were familiarized with the task with ∼10 practice trials.

### Behavioural Data Analysis

To assess the performance in the task, we first focused on the point of subjective equality (PSE), reflecting the accuracy of numerical estimates, and the just noticeable difference (JND) and Weber’s fraction (WF), reflecting the precision in the task. To compute these values, a cumulative Gaussian (psychometric) function was fitted to the proportion of “more numerous” responses as a function of the different levels of average numerosity, collapsing together the different numbers of arrays. The psychometric fitting was performed following the maximum likelihood method described by Watson, (1979). From the psychometric fit, we computed the PSE as the average numerosity corresponding to chance level responses, reflecting the perceptual match with the memorized reference. The JND was instead computed from the slope of the fit. As an additional measure of precision in the task, we computed the Weber’s Fraction (WF), which is the ratio of the JND and the PSE. This additional measure allows to assess the precision in the task while accounting for changes in the perceived magnitude of the stimuli. In addition to computing the general measures of performance, we also computed the accuracy (PSE) and precision (WF) of average numerosity judgments as a function of the number of arrays included in each sequence. To do so, we performed the psychometric fit separately for the trials in which the number of arrays was 3, 4, 6, 9 or 12. The PSE and WF were computed from these fits as explained above. To assess the biases in perceived average numerosity as a function of sequence length, we used a linear mixed-effect (LME) model test on the PSEs, entering the sequence length as predictor and the subject as the random effect.

In order to assess the weights of different arrays in the sequence in driving the judgment of average numerosity, we employed a non-linear regression analysis. The analysis was performed separately according to the number of arrays in the sequences, in order to further assess whether the amount of information provided affects the weighting profile of different arrays in the sequence. In the analysis, the binary response of the classification task was entered as the dependent variable, and the numerosity of the arrays along the sequence as the predictors. In order to have the same number of parameters across the different tests (i.e., to make the results more easily comparable across different tests), the predictors included only the first array in the sequence, the last array, and either the middle array (in the case of 3-array sequences) or an average of two intermediate positions in the sequence (second and third, third and fourth, fourth and sixth, and sixth and seventh, respectively for sequences of 4, 6, 9, and 12 arrays). The resulting beta values were first analysed using an LME model including serial position and sequence length as predictors (and subjects as the random effect) to assess the difference across the temporal weighting profiles as a function of the number of arrays in the sequence. Then, follow-up LME tests were performed within each level of sequence length.

Finally, we assessed the perceptual adaptation effects across successive trials, measuring how the perceived average numerosity of the stimuli is affected by the numerosity presented in the preceding trial. We employed again a psychometric fitting procedure, performed separately according to the average numerosity of the preceding stimulus. This analysis was repeated considering all the duration of the preceding stimulus together, and by separating the trials according to whether the preceding stimulus was short (3-4 arrays; 150-200 ms) or long (9-12 arrays; 450-600 ms). The PSEs obtained in this analysis were then used to compute an adaptation effect index, based on the difference in PSE between trials in which the preceding stimulus had a numerosity of 30 (the same as the reference; “PSE_30_”), and either lower (15, 21 dots) or higher (45, 60 dots) average numerosities ("PSE_j_”), according to the following formula:

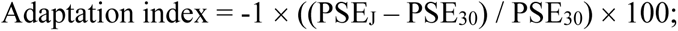

The sign was switched in order to make the interpretation of the index more intuitive, i.e., a negative adaptation index indicates a relative underestimation of the stimulus, while a positive index indicates an overestimation. The adaptation indexes were then analysed using LME models to assess the extent to which the perceived average numerosity of the current stimulus is affected by the stimulus in the previous trial.

### Electrophysiological recording and pre-processing

Throughout the experimental session we recorded the EEG in order to address the neural signature of average numerosity processing and the signature of the adaptation effect. The EEG was recorded using the Biosemi ActiveTwo system (2048 Hz sampling rate) and a 64-channel cap based on the 10-20 system layout. In order to more easily monitor artifacts due to eye movements and blinks, the electro-oculogram (EOG) was measured via an additional electrode attached below the left eye of the subject. The electrode offset values across the channels were usually kept below 20 µV, but occasional values up to 30 µV were tolerated.

The data pre-processing was performed offline in Matlab (version R2021b), using the functions of the EEGLAB (Delorme & Makeig, 2004) and ERPlab (Lopez-Calderon & Luck, 2014) toolbox. First, EEG signals were re-sampled to a sampling rate of 1000 Hz. Then, each combination of average numerosity and number of arrays was binned individually, for a total of 25 bins. Additionally, we added bins corresponding to the combination of different numerosities and different numbers of arrays of the stimuli in the previous trial, and different numerosities in the current trial, in order to assess the signature of adaptation effects (125 unique combinations). The continuous EEG data was then epoched time-locking the signal to the onset of each stimulus (i.e., the onset of the first array in each stimulus sequence). The epochs spanned from -300 ms to 1200 ms around the stimulus onset. The pre-stimulus interval (-300:0 ms) was used for baseline correction. The EEG signal was band-pass filtered with cut-offs at 0.1 and 40 Hz. To reduce artefactual activity in the data, we used an independent component analysis (ICA), aimed at removing identifiable artifacts such as eye movements and blinks. We additionally employed a step-like artifact rejection procedure (amplitude threshold = 40 μV, window = 400 ms, step = 20 ms) to further remove any remaining large artifact from the signal, leading to the exclusion of 2.9% ± 2.8% of the trials, on average (± SD). Finally, the event-related potentials (ERPs) were computed by averaging EEG epochs within each bin. ERPs were further low-pass filtered with a cut-off at 30 Hz, and smoothed with a sliding-window average with a width of 20 ms and a step of 5 ms.

### Event-related potentials analysis

The analysis of ERPs was performed by first selecting a set of channels of interest, based on previous studies. Namely, we selected a series of four occipital channels, including O1, O2, Oz, and Iz, based on previous studies on numerosity perception (Fornaciai et al., 2017; Fornaciai & Park, 2018) and trial-history effects in magnitude perception (Fornaciai et al., 2023; Tonoyan et al., 2022). First, we assessed numerosity-sensitive brain responses by sorting the ERPs according to the average numerosity of the stimuli, collapsing together the different durations. To assess the modulation of ERPs as a function of numerosity, we computed the linear contrast of the brainwaves (weights = [-2 -1 0 1 2], corresponding to the different levels of average numerosity). We then performed a series of one-sample t-tests against zero, corrected for multiple comparisons with a false discovery rate (FDR) procedure (q = 0.05). Moreover, we assessed the relationship between ERPs, in terms of the linear contrast amplitude, and behaviour in terms of the accuracy (PSE) and precision (JND) of average numerosity perception. To do so, we employed an LME model entering the contrast amplitude as the dependent variable, PSE and JND as predictors, and the subject as the random effect. To control for multiple comparisons, in this case we used a non-parametric cluster-based test. Namely, we repeated the analysis across the clusters of consecutive significant time windows observed in the actual LME test, randomly shuffling the vectors of PSE and JND values at each iteration. This procedure was repeated 10,000 times, and we measured how many times we could observe similar clusters of consecutive significant time windows in this simulation. The threshold used to consider a cluster significant in this analysis was conservatively set as the lower t-value observed in the corresponding cluster of the actual analysis, multiplied by the length of the cluster.

To assess the impact of adaptation effects on ERPs, we further sorted the data according to the average numerosity and number of arrays of the preceding stimulus, taking only the trials in which the middle numerosity (30 dots) was presented in the current trial. The modulation of ERPs was then tested by performing a series of LME tests across a series of small windows throughout the epoch, in a sliding-window fashion (width = 50 ms, step = 5 ms). The tests included the ERP amplitude as dependent variable, and the average numerosity and number of arrays (as well as their interaction) of the preceding stimulus. The subjects were added as the random effect. As the behavioural results showed an interaction between the numerosity and number of arrays of the previous stimulus, also in this case we considered an interaction between the factors as the crucial evidence for a signature of adaptation. To assess the nature of such an interaction, we further computed the effect of the preceding numerosity on ERP amplitude (i.e., the difference in amplitude corresponding to trials in which the preceding stimulus had 60 dots and 15 dots), as a function of the different number of arrays of the previous stimulus. This analysis was limited to the average ERPs within the latency window showing a significant interaction between average numerosity and number of arrays of the preceding stimulus. Again, to control for multiple comparisons we used a non-parametric cluster-based analysis (see above).

Finally, we assessed the relationship between the bias in perceived numerosity due to adaptation, and the extent to which adaptation modulates the ERP amplitude. To do so, we computed two corresponding measures of the adaptation effect on behaviour ("ι1PSE”) and on ERPs ("ι1ERP”). The ι1PSE was computed as the difference in PSE between the cases where the preceding stimulus had a numerosity either lower (15, 21 dots) or higher (45, 60 dots) than the reference (30 dots), and the case where the numerosity of the preceding stimulus was equal to the reference. The ι1ERP measure was computed in a similar fashion, as a difference in ERP amplitude in the cases where the preceding stimulus had either a lower or higher numerosity than the reference, and the case where the preceding stimulus had the same numerosity as the reference. To compute this index, we considered only the trials in which the stimulus in the current trial had 30 dots. We then used an LME model to assess the relationship between ΔPSE and ΔERP (ΔPSE ∼ ΔERP + (1|subj)). Again, this analysis was restricted to the latency window where the effect of adaptation on ERPs showed a significant interaction between the average numerosity and the number of arrays of the previous stimulus.

## RESULTS

In the present study we addressed the mechanisms of average numerosity perception and its neural signature. To do so, we used dynamic stimuli modulated in average numerosity (15-60 dots) and in the number of arrays composing the sequence (3-12, corresponding to a stimulus duration of 150-600 ms). Each array in the sequence could have a numerosity varying ±50% around the average. Although divided into briefly-presented (50 ms) but discrete arrays, the stimuli were designed to give the impression of a more continuous change. Indeed, with such a fast modulation of numerosity (20 Hz), the stimuli appeared like continuous stream of dots rather than a series of discrete frames. In the experiment, the participants (N = 21) performed a classification task, indicating whether the stimulus in each trial was more or less numerous, on average, compared to a memorised reference (average numerosity = 30 dots, 6 arrays), presented before each block of trials. See Fig. 1 for a depiction of the experimental procedure.

**Figure 1.**
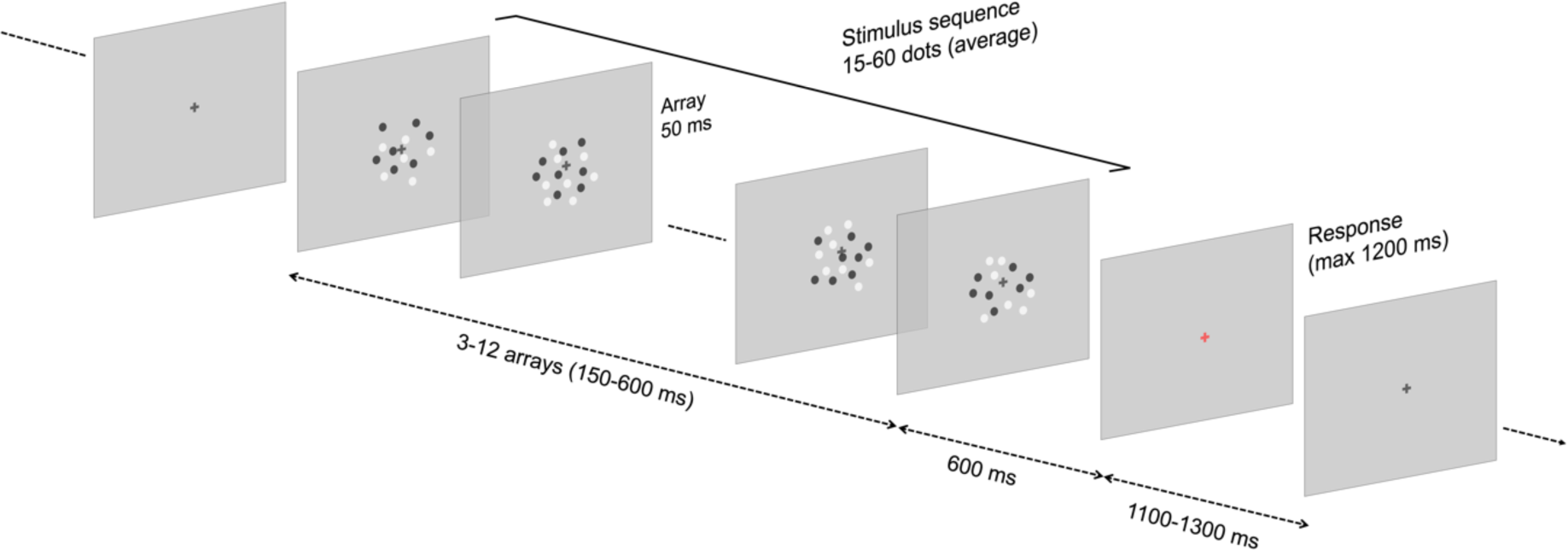
Stimulation procedure. The classification task involved participants watching a series of dynamic stimuli modulated in average numerosity and in the number of arrays presented, and determining whether the average numerosity in each trial was higher or lower compared to a memorized reference stimulus. The reference was presented at the beginning of the session and at the beginning of each block. Each array in the stimulus sequence included a set of black and white dots drawn within a circular area, presented for 50 ms. The number of arrays presented in each sequence varied from 3 (150 ms) to 12 (600 ms). Even if composed of individual arrays, the stimulus was designed to appear as a continuous stream rather than a series of discrete stimuli. The offset of the stimulus sequence was followed by a 600 ms blank interval. After the interval, the fixation cross became red, signalling to the participant to provide a response by pressing the appropriate key on a standard keyboard. The next trial started automatically after an inter-trial interval of 1100 to 1300 ms. The stimuli are not depicted in scale.

First, we assessed the general performance of average numerosity judgments, computing measures of accuracy (point of subjective equality, PSE), and precision (just noticeable difference, JND, and Weber’s fraction, WF). Fig. 2A shows the general measures of performance. On average, the perceived numerosity of the stimuli was quite accurate and precise, showing a PSE (± SD) of 31.42 ± 4.41 dots, a JND of 7.70 ± 2.34 dots, and a WF of 0.25 ± 0.11. These results show that participants are able to judge average numerosity fairly accurately and precisely, in line with previous studies (Katzin et al., 2021; Togoli et al., 2021).

**Figure 2.**
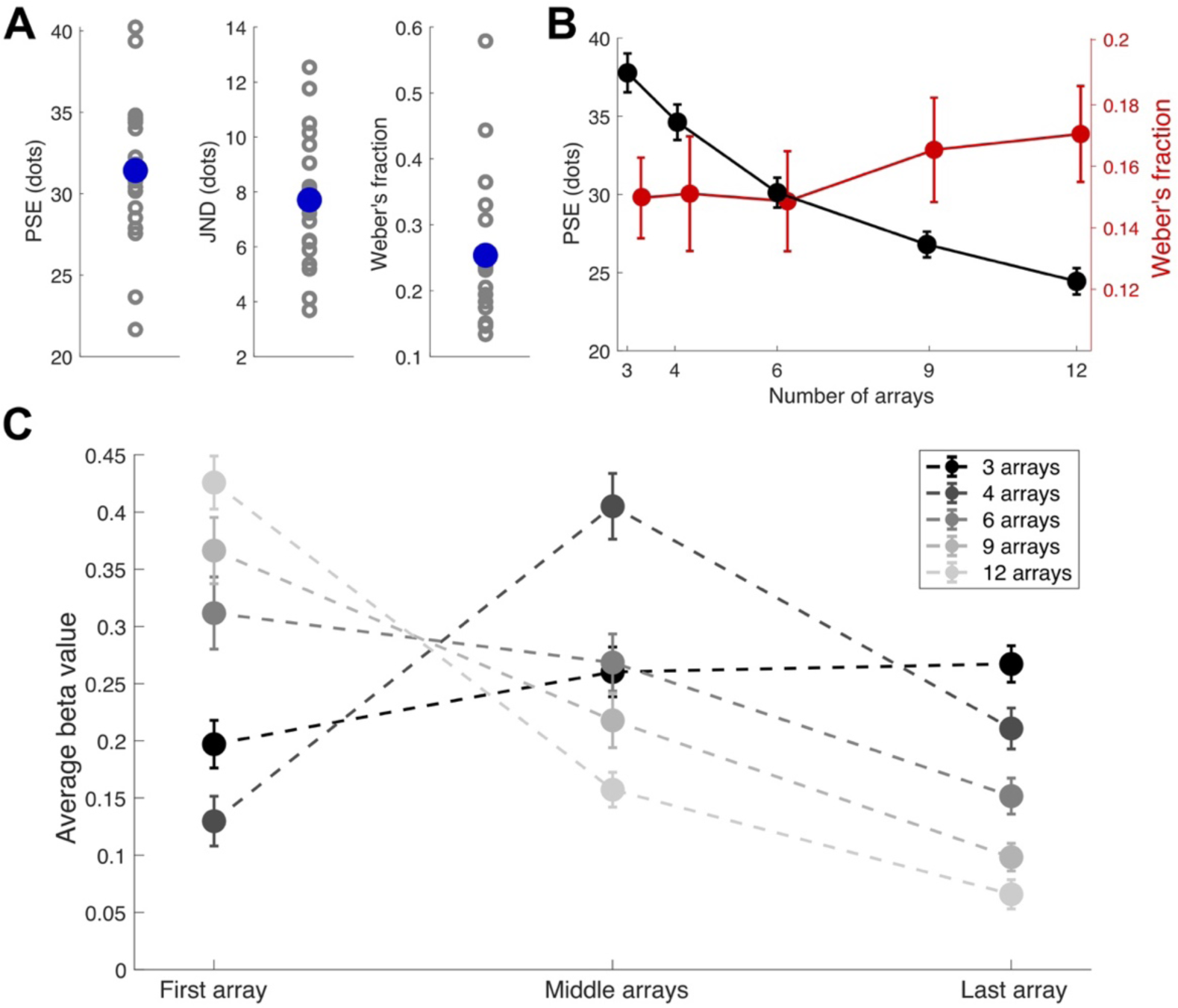
Behavioural results. (A) General measures of performance, including the point of subjective equality (PSE) as a measure of accuracy, and the just noticeable difference (JND) and Weber’s fraction (WF) as a measure of precision. (B) Biases in perceived average numerosity as a function of the number of arrays presented, and modulation of the precision of judgments (WF). (C) Temporal weighting profile, reflecting the weights of the first, middle, and last array in the sequence in driving the judgement, separately for the different numbers of arrays composing the sequence. Error bars are SEM.

First, an interesting question is whether the amount of information provided, i.e., the number of arrays presented in the sequence, affects the accuracy and/or the precision of judgments. To address this question, we computed the PSE and WF separately as a function of the number of arrays of the sequences (Fig. 2B). The results showed substantial biases in perceived numerosity according to the length of the sequence, with a pattern of both under- and over-estimation. Indeed, when the number of arrays was smaller than the reference, PSEs were higher (37.78 ± 1.24 and 34.62 ± 1.13, respectively for 3 and 4 arrays), showing an underestimation of perceived average numerosity (i.e., a higher number of dots is necessary to perceptually match the memorised reference). Conversely, PSEs were lower when the sequence was longer than the reference (26.80 ± 0.82 and 24.44 ± 0.84, respectively for 9 and 12 arrays), showing an overestimation. A linear mixed-effect regression model (LME; PSE ∼ Number of arrays + (1|subj)) showed a significant modulation of PSE as a function of sequence length (adj-R^2^ = 0.85, β = -1.43, t = -17.69, p < 0.001). On the other hand, the WF (shown in red in Fig. 2B) showed a slight but significant increase with sequence length (LME test on WFs; adj-R^2^ = 0.76, β = 0.002, t = -2.27, p = 0.025). This suggests that precision was lower when the sequence included more arrays than the reference.

Furthermore, we addressed the temporal weighting profile of the dynamic stimuli – that is, the extent to which arrays in different positions along the sequence contribute to the perception and judgment of average numerosity. To do so, we employed a non-linear regression analysis, entering the binary classification response as the dependent variable, and the numerosity of arrays along the sequence as predictors. This analysis was performed separately as a function of the number of arrays in the sequence, to further assess whether the amount of information presented affected the weight of different portions of the sequence. To ensure that the results concerning different sequences are comparable, we included the same number of parameters (array position) in the non-linear regression: the first and the last array, and either the middle array (in the case of 3 arrays) or the average of two intermediate arrays. The beta values obtained with this analysis reflect the extent to which arrays in different positions contribute to the classification judgement.

The results, shown in Fig. 2C, show a clear difference in the temporal weighting profile as a function of the sequence length. A linear mixed effect (LME) test on the beta values, with factors position and number of arrays, showed indeed a significant interaction across the two factors (adj-R^2^ = 0.37, β = -0.025, t = -10.93, p < 0.001). Interestingly, considering the pattern across the three sequence positions (first, middle, last; Fig. 2C), the change in beta values showed different directions according to the number of arrays presented. Follow up LME tests on beta values performed individually within each level of sequence length (i.e., 3-12 arrays) first showed a significant difference in the 3-array sequence, in the positive direction, i.e., higher beta values at the end of the sequence (adj-R^2^ = 0.08, β = 0.035, t = 2.55, p = 0.013). With 4 arrays, the middle position showed a higher weight, but overall the difference was not significant (adj-R^2^ = 0.03, β = 0.040, t = 1.73, p = 0.09). Conversely, with sequences longer than 4 arrays, the tests showed significant differences in the negative direction, reflecting a higher weight of the first array in the sequence compared to the last (adj-R^2^ = 0.24, β = -0.080, t = -4.59, p < 0.001, adj-R^2^ = 0.53, β = -0.134, t = -8.52, p < 0.001, and adj-R^2^ = 0.73, β = -0.180, t = -13.09, p < 0.001, respectively for 6, 9, and 12 arrays). This suggests that the weighting of different arrays in the sequence is subject to either a recency effect (i.e., larger weight of information presented at the end of the sequence; see for instance Hubert-Wallander & Boynton, 2015) or a primacy effect (i.e., larger weight to information presented earlier in the sequence), depending on the length of the sequence itself.

An additional aspect that we assessed is the possibility of perceptual adaptation effects across stimuli in different trials. Indeed, even if very brief (150-600 ms), the stimuli might potentially induce perceptual adaptation effect, as sequences of brief arrays have been previously shown to induce significant effects (Aagten-Murphy & Burr, 2016). If present, adaptation effects might provide more insights into the representation of perceived average numerosity. To address this possibility, we computed the perceived average numerosity of the stimuli as a function of the average numerosity preceding them. The results of this analysis are shown in Fig 3. First, we observed a robust modulation of the perceived numerosity of the stimuli (PSE) as a function of the preceding numerosity (Fig. 3A). To better assess the bias induced by the previous stimuli, we computed an adaptation effect index reflecting how much the perception of the stimuli changes when the previous stimulus had either a lower or higher numerosity than the intermediate, reference numerosity (Fig. 3B). In line with the repulsive nature of perceptual adaptation, we observed a relative overestimation when the preceding stimulus had fewer dots than the reference (15, 21 dots), and a relative underestimation when the preceding stimulus had more dots (45, 60 dots). The biases ranged from 6% to about -8%. An LME test showed a significant difference in the adaptation effect as a function of the preceding numerosity (adj-R^2^ = 0.61, β = -0.306, t = -6.37, p < 0.001). Moreover, a perceptual adaptation effect can also be expected to increase with either the duration or the number of arrays presented in the previous stimulus (Aagten-Murphy & Burr, 2016). We thus further tested whether the number of arrays could modulate the effect. The PSEs shown in Fig. 3C indeed suggest that when the preceding stimulus was longer (9, 12 arrays), the bias in perceived numerosity was stronger compared to a shorter sequence (3, 4 arrays), especially at larger numerosities. To assess the pattern of adaptation effects computed as a function of the number of arrays of the preceding stimulus (Fig. 3D), we used an LME model test with factors numerosity and number of arrays of the preceding stimulus. The results showed a significant interaction between the two factors (adj-R^2^ = 0.60, β = -0.271, t = -2.44, p = 0.016), suggesting a stronger effect when the preceding sequence was longer, especially at larger numerosities. Further LME tests, performed individually for the two sequence lengths, showed however that in both cases the effect is statistically significant (adj-R^2^ = 0.71, β = - 0.179, t = -3.51, p < 0.001, and adj-R^2^ = 0.70, β = -0.450, t = -5.53, p < 0.001, respectively for short and long sequences).

**Figure 3.**
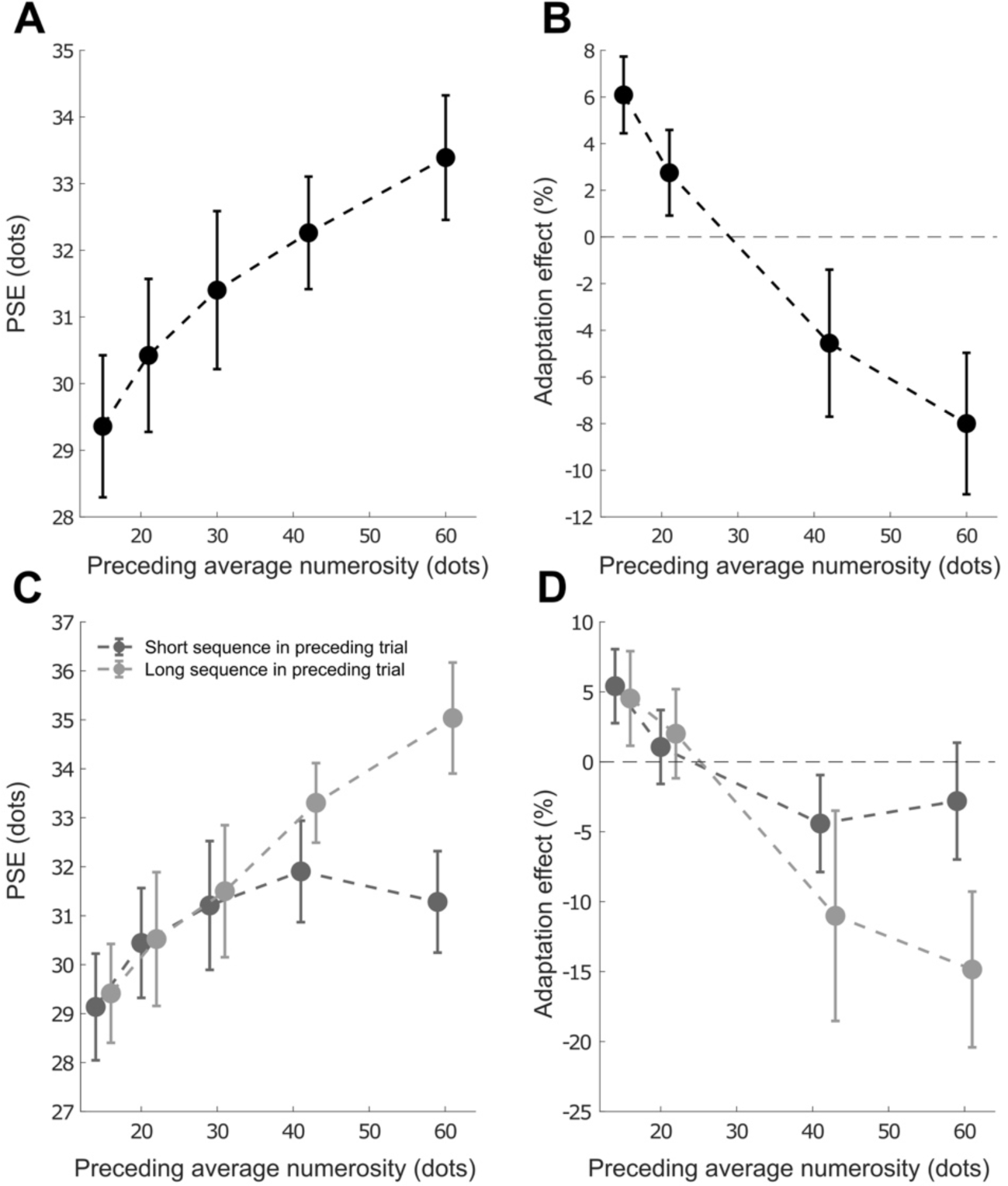
Perceptual adaptation effects. (A) PSEs as a function of the preceding numerosity. (B) Average adaptation effect indexes as a function of the preceding numerosity. (C) PSEs as a function of the preceding numerosity, separately for the cases where the previous stimulus had a short (3, 4 arrays) or long (9, 12 arrays) sequence. (D) Adaptation effect indexes as a function of preceding numerosity and sequence length (short vs. long). Error bars are SEM.

After characterising the properties of average numerosity perception at the behavioural level, we addressed the neural signature of this process. First, we assessed the brain responses sensitive to average numerosity, which are shown in Fig. 4A. To do so, we sorted the ERPs according to corresponding average numerosity, and computed the linear contrast of ERPs as a measure of sensitivity to numerical information. The linear contrast was then tested with a series of one-sample t-tests against zero, corrected for multiple comparisons with a false discovery rate (FDR) procedure (q = 0.05). The results of the series of tests showed numerosity-sensitive activity in four distinct latency windows, starting very early after the onset of the sequence: 20-55 ms (t(20) ≥ 3.10, adjusted-p ≤ 0.039), 115-155 ms (t(20) ≤ -3.35, adj-p ≤ 0.026), 190-230 ms (t(20) ≥ 2.99, adj-p ≤ 0.049), and 430-595 ms (t(20) ≤ -2.98, adj-p ≤ 0.049). This shows that brain responses to the stimulus can track the numerosity of the sequence starting shortly after its onset.

**Figure 4.**
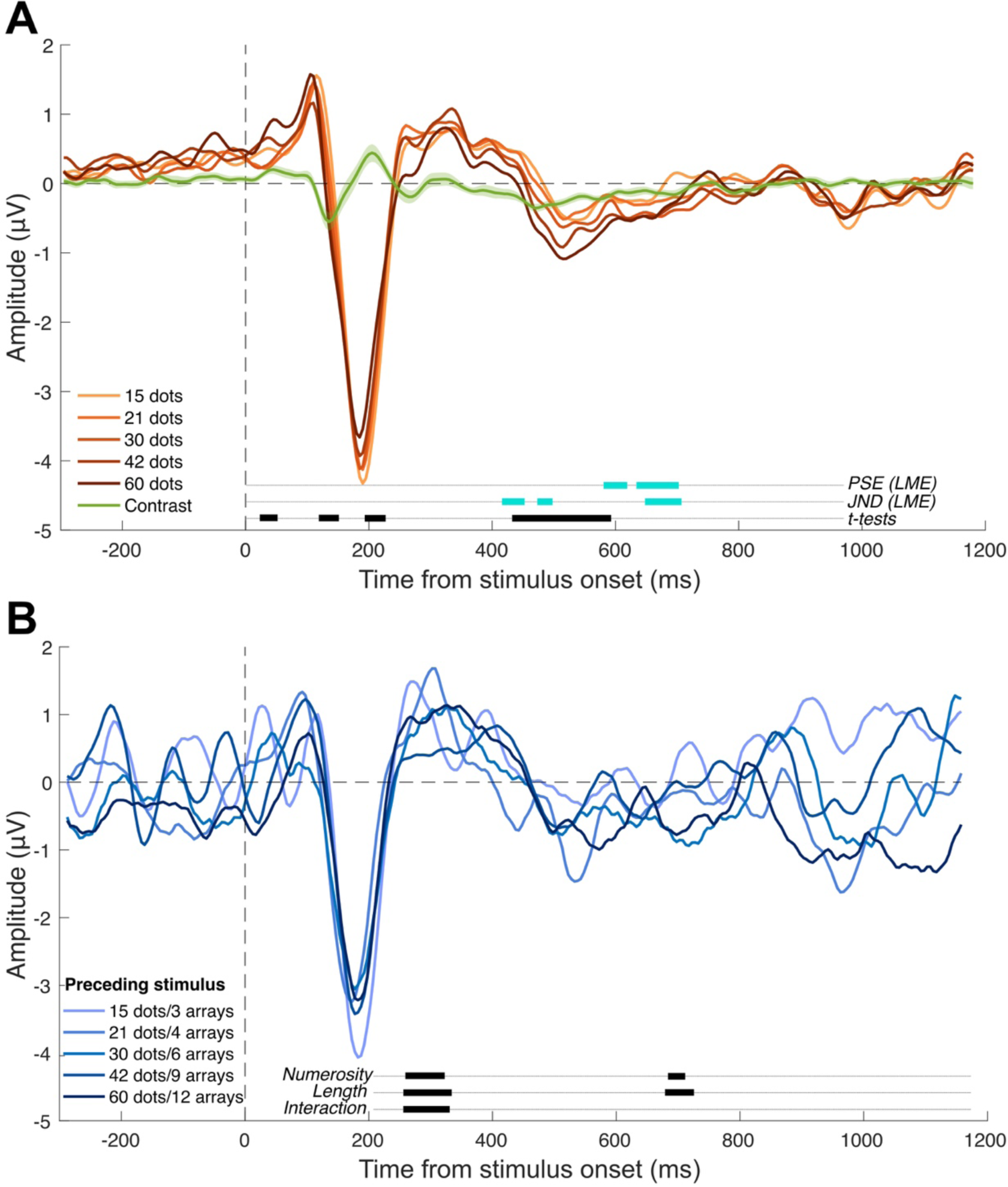
Event-related potentials (ERPs) reflecting average numerosity and adaptation effects. (A) ERPs sorted according to average numerosity. The green wave represents the linear contrast of ERPs. The shaded area represents the SEM. Lines at the bottom of the plot show the significance of different tests. The thick black lines indicate the latency windows where the linear contrast is significantly different from zero (FDR-corrected one-sample t-tests). The cyan lines instead indicate the results of the LME test relating the ERPs to the behavioural performance. Namely, the lines show the latency windows where we observed a significant relationship between the amplitude of the linear contrast and either the PSE or the JND. (B) Representative ERPs reflecting the effect of the preceding stimulus on the responses to the intermediate average numerosity in the current trial (30 dots). For the sake of clarity, only the pairwise corresponding combinations of average numerosity and number of arrays (length) are shown in the plot. However, the analysis was performed on the full set of 25 unique combinations of average numerosity and length of the preceding stimulus. The lines at the bottom of the plot show the results of the LME tests, reflecting the significance of the average numerosity, length, and their interaction in driving the ERPs.

However, a question in this context is: at what processing stage does the average numerosity is actually computed? And what are the processing stages contributing to the judgement of average numerosity? To answer these questions, we focused on the relationship between brain activity and behavioural performance, in terms of the accuracy (PSE) and precision (JND) of numerical judgments. We thus performed LME tests on the contrast amplitude throughout the post-stimulus interval, entering PSE and JND as predictors and the subjects as the random effect. The results first showed that the amplitude of numerosity-sensitive responses can be predicted by the PSE at two latency windows: 580-625 ms (adj-R^2^ = 0.62-0.65, β ≤ -0.022, t ≤ -2.11, p ≤ 0.049), and 635-710 ms (adj-R^2^ = 0.61-0.87, β ≤ -0.026, t ≤ -2.13, p ≤ 0.047).

Both these windows show a negative relationship between amplitude and PSE, suggesting that higher amplitudes are associated with lower PSE. Second, the results also showed a significant relationship between the contrast amplitude and JND at three latency windows: 410-455 ms (adj-R^2^ = 0.62-0.65, β ≥ 0.045, t ≥ 2.22, p ≤ 0.039), 470-500 ms (adj-R^2^ = 0.63-0.67, β ≥ 0.051, t ≥ 2.36, p ≤ 0.030), and 650-715 ms (adj-R^2^ = 0.66-0.86, β ≤ -0.045, t ≤ -2.25, p ≤ 0.037). Interestingly, both earlier windows show a positive relationship between amplitude and JND, suggesting that higher amplitudes are associated with poorer precision. Instead, at the later window (650-715 ms), the negative relationship suggests that the higher the sensitivity of responses to numerosity, the higher the precision (i.e., the lower the JND). These results show that while numerosity-sensitive activity emerges throughout the presentation of the stimulus sequence, only later activity show a relationship with the judgment of average numerosity. To control for multiple comparisons, the significant windows observed in the LME tests were assessed with a non-parametric cluster-based test (see Methods). All the cluster p-values resulted to be < 0.001.

Since we observed significant adaptation effects at the behavioural level, addressing the neural signature of such effects can provide further insights into the processes contributing to the perception of average numerosity. In this context, we took the ERPs corresponding to the presentation of 30 dots in the current trial, and sorted them according to the average numerosity and sequence length of the preceding stimuli. Fig. 4B shows a representative set of the ERPs reflecting the responses to the “current” (i.e., average numerosity of 30 dots) stimulus as a function of the preceding stimulus. In the analysis, however, we considered the full set of 25 unique combinations of average numerosity and number of arrays of the preceding stimulus. We then performed LME tests on the ERP amplitude throughout the post-stimulus interval, adding the numerosity and the sequence length of the preceding stimulus as predictors, and the subjects as the random effect. Since we observed an interaction between numerosity and length in the behavioural effect of adaptation, our prediction is that a signature of the effect should show a similar interaction between the two factors. The results first show a significant effect of numerosity on the ERP amplitude at two latency windows: 245-330 ms (adj-R^2^ = 0.55-0.66, β ≤ -0.025, t ≤ -2.33, p ≤ 0.020) and 670-725 ms (adj-R^2^ = 0.14-0.15, β ≤ -0.021, t ≤ -1.97, p ≤ 0.049). The effect of sequence length was observed at two similar latency windows: 240-350 ms (adj-R^2^ = 0.51-0.67, β ≤ -0.002, t ≤ -1.97, p ≤ 0.048) and 665-735 ms (adj-R^2^ = 0.12-0.16, β ≤ -0.002, t ≤ -2.07, p ≤ 0.039). Additional significant effects of the sequence length were observed at other latencies where no other effect was found. Since an effect of the preceding sequence length in isolation (i.e., not coupled with an effect of numerosity or an interaction) is difficult to interpret in this context, we did not consider such latency windows for further analysis. More importantly, we observed a significant interaction between numerosity and sequence length at 240-345 ms (adj-R^2^ = 0.51-0.67, β ≥ 0.001, t ≥ 2.04, p ≤ 0.042). These significant latency windows were further assessed with non-parametric cluster-based tests to control for multiple comparisons. All the cluster p-values were < 0.001.

To better understand the nature of this interaction, we assessed how the effect that numerosity plays through adaptation is modulated by the length of the sequence, within the latency window showing a significant interaction between the two factors (i.e., average of amplitudes within 240-345 ms after stimulus onset). We thus computed a measure of the effect of numerosity in the preceding trial on ERPs (i.e., amplitude of responses to 60 dots minus responses to 15 dots), and plotted it as a function of the length of the preceding stimulus (Fig. 5A). The results show indeed a clear modulation, with a significant effect of the preceding stimulus length in modulating the impact that numerosity has on ERPs (LME test; adj-R^2^ = 0.08, β = 0.193, t = 3.14, p = 0.002). This test shows however a low R2, which may be due to the difficulty of the linear model to capture a potentially non-linear effect (see the pattern in Fig. 5A).

**Figure 5.**
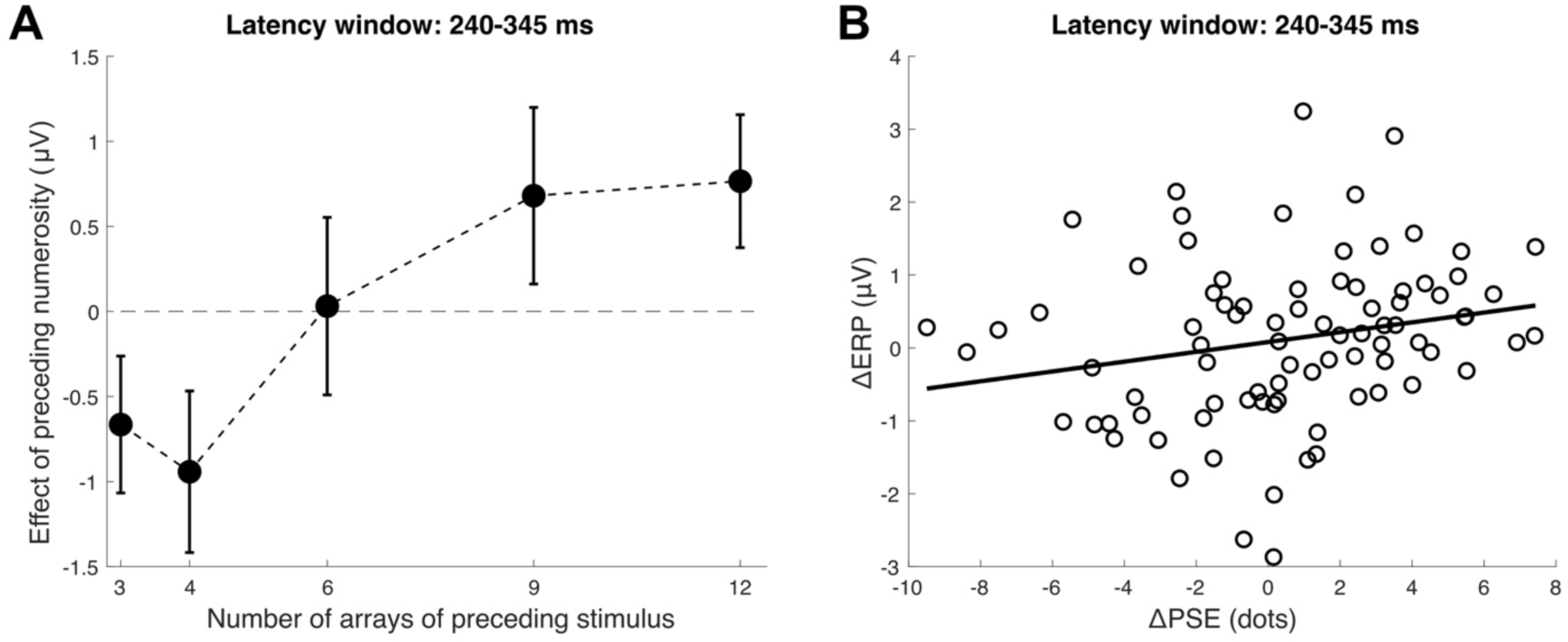
Interaction between numerosity and length and relationship between the behavioural and electrophysiological effect of adaptation. (A) Effect of the preceding numerosity as a function of the length of the preceding sequence. Error bars are SEM. (B) Effect of adaptation measured behaviourally (11PSE) as a function of the effect measured with EEG (11ERP). The 11PSE and 11ERP measures were computed as the difference in PSE or ERP amplitude between either lower (15, 21 dots) or higher (45, 60 dots) numerosities of the preceding stimulus and the intermediate numerosity of the range (30 dots). The line represents a linear fit to the data, to show the general trend. The data was however analysed with an LME model. Both panels report data averaged within the latency window where we observed a significant interaction between the numerosity and length of the preceding stimulus (240-345 ms after stimulus onset).

Finally, we assessed the relationship between the behavioural and electrophysiological signature of the adaptation effect. To do so, we computed two corresponding measures: 11PSE and 11ERP. These measures were computed as the difference in either PSE or ERP amplitude between the lower (15, 21 dots) and higher (45, 60 dots) numerosity levels of the previous stimulus and the intermediate level (30 dots). The ΔERP was computed in a similar fashion, but considering an average numerosity of 30 dots in the current stimulus. This measure was computed as the average within the latency window showing an interaction between numerosity and length of the preceding stimulus (240-345 ms). Fig. 5B shows the general trend of these data. To assess the relationship between the two measures, we performed an LME test, entering the ΔPSE as the dependent variable, and 11ERP as the predictor, and adding the subject as the random effect. The results showed a significant relationship, whereby the ΔPSE, indexing the behavioural effect of adaptation, can be predicted by the ΔERP, which indexes the changes in ERP amplitude due to the preceding stimulus (adj-R^2^ = 0.56, β = 0.813, t = 2.26, p = 0.026). The relationship is positive, suggesting that the higher the change in ERP amplitude within the 240-345 ms window, the higher the behavioural effect of adaptation.

## DISCUSSION

In the present study, we addressed the mechanisms of average numerosity perception with fast dynamic stimuli, and the neural signature of this process. How the brain computes and represents the approximate average of numerosity over time indeed remains mostly unclear. So far, only a few studies addressed this process, providing initial evidence for the existence of dedicated brain mechanisms supporting the averaging of numerosity information over time (Katzin et al., 2021; Togoli et al., 2021). Our results provide new evidence showing that (1) the weighting profile of information along the sequence depends on the total amount of information provided; (2) average numerosity is subject to perceptual adaptation effects across trials; and (3) average numerosity and the adaptation effect show robust neural (electrophysiological) signatures, with activity at specific latency windows predicting the behavioural performance and effects.

First, in terms of general performance, we observed a few differences compared to previous results. Namely, while Katzin et al. (2021) reported that the averaging precision increases as a function of the sequence length, here we observed a slight but significant reduction in precision (i.e., Weber’s fraction). Additionally, we also observed biases in accuracy depending on the sequence length, with both under- and over-estimation of average numerosity according to whether the sequence was shorter or longer than the reference, respectively. These biases are in line with the influence of duration on perceived numerosity (e.g., Javadi & Aichelburg, 2012; Lambrechts et al., 2013; Walsh, 2003), and suggest that our dynamic stimuli were indeed perceived as a continuous sequence rather than a series of discrete stimuli. Indeed, it was the entire duration of the sequence – not the duration of individual arrays (which was constant) – that affected the overall perceived average numerosity. Recent results also show that using dynamic stimuli amplifies the effect of duration on numerosity (Togoli et al., 2022), and the strength of the bias observed here is consistent with such previous results. When it comes to the difference between our results and those of Katzin et al. (2021), a possible explanation is a difference in the nature of the stimuli used. Namely, while Katzin et al. used sequence of relatively long stimuli (500 ms), which are most likely perceived as different, discrete arrays, here we used a fast modulation of numerosity (20 Hz) that is likely perceived as a single continuous stimulus. The difference in how the sequence length modulates accuracy and precision may thus suggest that computing the average of continuous vs. discrete sequences may engage different mechanisms.

Despite the relatively short duration of the stimulus sequences, we observed systematic perceptual adaptation effects. Numerosity adaptation effects are usually observed with much longer exposures, around a few seconds (e.g., Arrighi et al., 2014; Burr & Ross, 2008; Grasso et al., 2022). However, there is also evidence that adaptation can be observed even with very short exposures, at least in the case of ambiguous or masked stimuli (Fornaciai & Park, 2019b, 2021; Glasser et al., 2011). Additionally, numerosity adaptation effects have been observed after repeated presentations of brief arrays of dots (Aagten-Murphy & Burr, 2016), making our effect in line with previous results.

More importantly, we observed different temporal weighting profiles according to the sequence length. While sequences up to four arrays (or 200 ms) showed either a recency effect or a flat profile, longer sequences showed primacy effects. This pattern is particularly interesting, as it suggests an important capacity limit for the computation of average numerosity. Namely, with a relatively short sequence, the visual system is able to use the information provided with similar weights, giving however more relevance to more recent information. When the sequence gets longer (6-12 arrays, 300-600 ms), the visual system weights further information increasingly less compared to the early information. Comparing these temporal weighting profiles with results obtained in other feature dimensions, the primacy effect observed with long sequences is similar to the one observed by Hubert-Wallander & Boynton (2015) on average spatial position. Hubert-Wallander & Boynton (2015) also tested and compared different dimensions, such as size, face identity, and motion, which mostly showed recency effects. Our results thus add to those of Hubert-Wallander & Boynton (2015) in suggesting the existence of different, dimension-specific mechanisms for the extraction of summary statistics.

What is the mechanism underlying the peculiar temporal weighting profiles observed in average numerosity? The weighting profile may be due to two possible types of capacity limits, either cognitive or perceptual. First, the capacity limitation leading to difference weighting profiles may be due to the limits of working memory (WM) encoding, usually considered to be around 3-4 items (Alvarez & Cavanagh, 2004; Awh et al., 2007; Luck & Vogel, 1997). This limit of WM encoding seems indeed consistent with our results, showing that information exceeding the third or fourth array in the sequence is given increasingly less weight. However, one might expect that with longer sequences more recent information would be encoded in working memory replacing older information, leading to a recency effect (as found by Katzin et al. 2021 with sequences of discrete stimuli). A recency effect was however observed only at the shorter sequence. As mentioned above, differently from previous studies (Katzin et al., 2021) our stimuli were designed to appear like a continuous stream rather than a sequence of discrete stimuli. A second, and perhaps more plausible explanation for the weighting profiles is thus a perceptual capacity limitation, based on the limits of temporal integration of the early visual system. Indeed, the ability of the visual system to integrate information during the presentation of a stimulus is limited by several factors, including for instance rapid adaptation of responses and the correlation in response fluctuations (e.g., see for instance Goris et al., 2018). Due to these constraints, the limit of temporal integration in visual cortex has been estimated to be around 150-300 ms (e.g., Burr & Santoro, 2001; Goris et al., 2018)). After this time, the temporal integration (or summation) of signals – and its benefits in visual perception – tend to plateau. The timing of our weighting profiles seems consistent with this time course, as the turning point after which the weights start to decrease is around 200 ms (4 arrays). A possibility is thus that the summary statistics of a relatively brief, continuous sequences may purely rely of the ability of the early visual system to track and integrate information over time, rather than by the encoding of information in visual working memory.

The EEG results provide further evidence for the mechanisms involved in the computation of average numerosity. First, when assessing numerosity-sensitive brain responses at occipital channels (Iz, Oz, O1, O2), we observed significant activity starting very early after stimulus onset (20-50 ms). Numerosity-sensitive responses continued throughout the presentation of the stimulus, with the more sustained activity emerging around the offset of the longer stimulus sequences (430-595 ms). The initial onset of such responses is extremely early in terms of latency, but it may be consistent with early responses to numerosity observed in previous studies (e.g., ∼75 ms in Park et al., 2016, 50-80 ms in Fornaciai et al., 2017; in both cases with signals measured at Oz). Such an early onset may thus be consistent with the responses of early visual areas such as V1, V2, and V3 (e.g., Fornaciai et al., 2017; Foxe & Simpson, 2002). To better understand how the brain responses evoked by the stimuli are related to the perception and judgment of average numerosity, we also assessed the relationship between the amplitude of ERPs and the behavioural measures of accuracy (PSE) and precision (JND). The results show a series of latency windows whereby the behavioural performance can predict the amplitude of ERPs, clustered around the offset of the longer stimulus sequences (∼400-700 ms). This suggests that these processing stages may be related to the actual computation of average numerosity based on responses integrated during the stimulus presentation. Another possibility, however, is that such late stage might represent a higher level perceptual decision-making stage involved with the judgement of average numerosity.

Moreover, the adaptation effect shows instead an earlier signature, with occipital responses at ∼240-340 ms reflecting the interaction between the average numerosity and the length of the preceding sequence. First, such a localised window suggests that adaptation did not affect the general visual responses to the stimuli or the perceived numerosity of the individual arrays in the sequence, but more likely the computation of summary statistics from the stimulus sequence. Second, activity within this latency window can predict the adaptation effect measured behaviourally. Thus, these aspects of the adaptation effect suggest that the observed latency window may reflect a processing stage involved with the computation of perceived average numerosity. Previous results concerning numerosity adaptation (Grasso et al., 2022) observed a correlate of the effect at the P2p component, an ERP component usually associated with numerosity processing (Fornaciai et al., 2017; Fornaciai & Park, 2018; Libertus et al., 2007; Park et al., 2016). The latency window observed here is not far off from the typical timing of the P2p (200-250 ms), but due to the different nature of the stimuli it may represent a different computational stage. A possibility is that such an intermediate stage might be more genuinely involved with the computation and representation of average numerosity compared later latency windows showing a relationship with accuracy and precision. Such later stages, as mentioned above, might instead reflect perceptual decision-making processes. Note however that all these different ERP analyses were performed considering the same set of occipital electrodes (Iz, Oz, O1, O2). Although the spatial resolution of EEG is notoriously poor, these results are nevertheless more consistent with perceptual processing occurring in visual cortex rather than cognitive processes occurring in higher-level brain areas.

To conclude, our results provide new insights into the computation of summary statistics in numerosity perception, demonstrating the properties and neural signature of average numerosity. Our results overall suggest the existence of specific low-level mechanisms dedicated to the computation of average numerosity over time, subject to the limits of temporal integration of early visual areas. The neural signature of average numerosity and the adaptation effect further show two crucial processing stages, at intermediate (∼300 ms) and late (∼600 ms) latencies. These stages are potentially consistent with the initial representation of average numerosity and a subsequent perceptual decision-making stage.

## Acknowledgements

This project has received funding from the European Union’s Horizon Europe research and innovation programme under the Marie Sklodowska-Curie grant agreement No. 101103020 “PreVis” to MF, from the European Research Council (ERC) under the European Union’s Horizon 2020 research and innovation programme grant agreement No. 682117 BIT-ERC-2015-CoG to DB, and from the Italian Ministry of University and Research under the call FARE (project ID: R16X32NALR) and under the call PRIN2017 (project ID: XBJN4F) to DB.

